# A cGAS-mediated IFN-I response in human CD4+ T cells depends on productive infection and is conserved over HIV types and strains

**DOI:** 10.1101/2024.01.12.575325

**Authors:** Marija Janevska, Timothy Cammaert, Evelien Naessens, Bruno Verhasselt

## Abstract

HIV type 2 is known to be better controlled by our immune system than HIV-1. The mechanism of innate sensing of HIV-2 by T cells is at present unclear. In this study we show that several primary isolates of HIV-2 (CBL20 and CI85) and HIV-1 (A8 and D2), similarly to the molecular clone HIV-1 NL4.3-GFP-I, induce a significant IFN-I response by infection of its main target, activated CD4+ T cells. However, they are unable to do so after shRNA-mediated knock-down of cGAS. In addition, HIV-1 induced IFN-I response in CD4+ T cells is dependent on productive infection and cannot be attributed to contaminating plasmid DNA present in some virus stocks. Our findings collectively showed that the cGAS-dependent innate response of CD4+ T cells to HIV infection is conserved over HIV types and critically depends on productive infection.

**Author Summary:** Our study unveils the essential role of cGAS in sensing HIV-1 and HIV-2 infections in CD4+ T cells. By demonstrating the necessity of productive infection, we highlight the robust and specific nature of the observed cGAS-mediated innate response, dispelling concerns about contaminating plasmids triggering immune response. Our findings suggest that the lower pathogenicity of HIV-2 does not correlate to superior innate immune control mediated by cGAS. By emphasizing the importance of productive infection and cGAS activity, our research advances understanding of host-pathogen relation and informs targeted strategies for combating HIV.

## Introduction

The slower progression and long-term asymptomatic presentation of HIV-2 infection raises questions about its ability to induce a potent interferon response when being sensed, and how it compares to HIV-1 in terms of signaling pathways involved. Both viruses (Human Immunodeficiency Virus (HIV) 1 and 2) have been introduced in the human population via different pathways. HIV-1 originates from Simian Immunodeficiency viruses (SIV) of the chimpanzee, while HIV-2 is known to come from SIV of the sooty mangabey [1–3]. This distinct origin and evolution is responsible for the distinct genetic build-up and diversity between HIV-1 and HIV-2 that can explain the vast differences between them [4].

Type-I interferons (IFNs) are triggered when pathogen-associated molecular patterns (PAMPs), like viral RNA, DNA or viral proteins are being sensed by different pattern recognition receptors (PRRs) expressed by infected cells. In order to contain the viral infection, the immune response of our body to viral pathogens involves induction of numerous IFN-stimulated genes (ISGs). This creates an “antiviral state” resulting in inhibition of viral replication.

An important PRR is the DNA sensor cGAS [5,6] which plays a crucial role in intracellular defense responses by sensing viral double-stranded DNA (dsDNA). Upon binding to viral DNA, cGAS catalyzes the production of cyclic GMP-AMP (cGAMP), which in turn activates the stimulator of interferon genes (STING) pathway, leading to the production IFNs. It has been demonstrated that HIV-1 infections triggers a cGAS-dependent immune response in macrophages [7–9] and activated CD4+ T cells [10], however evidence for HIV-2 is lacking.

Even though inhibitors of viral replication could block HIV-1 sensing, some authors argued the importance of cGAS was an artefact due to contaminating plasmid in the virus stocks used in some experiments [11].

Here, we investigated innate sensing of productive HIV infection of its main target, activated CD4+ T cells and the role of the DNA sensor cGAS. Our results show that innate sensing of HIV-2 occurs during productive infection, similarly to HIV-1, dependent on cGAS. This dependency was not explained by contaminating plasmid DNA in viral stocks, as upregulation of ISGs was observed in infection with cell-grown virus from primary isolates; heat sensitive and not diminished by DNA digestion.

## Results

### Innate response of CD4+ T cells to HIV infection is not an artefact caused by contaminating plasmid in virus stocks

We previously showed [10] that activated CD4+ T cells sense productive HIV-1 infection through the DNA sensor cGAS and induce an IFN-I response, that could be blocked by inhibitors of viral replication. However, it was argued that when using viruses produced by transfection of proviral plasmid DNA, the innate response could be attributed to the cGAS-depended sensing of plasmid DNA and not to sensing of HIV-1 infection [11]. In their paper, Elsner etal. used R5-tropic HIV-1 BaL strain to infect activated CD4+ T cells which failed to induce cGAS-dependant type I interferon response.

To show directly that plasmid contamination is not responsible for IFN induction during HIV infection in activated CD4+ T cells in our experiments, we first used heat-inactivated viral supernatant. As shown in Fig. 1A, after heat treatment of HIV-1 NL4-3 GFP-I supernatant, no infection nor IFN-I induction in IL-2/PHA–stimulated primary CD4+ T cells was observed, arguing against a role for heat-stable components like plasmids.

**Fig. 1.**
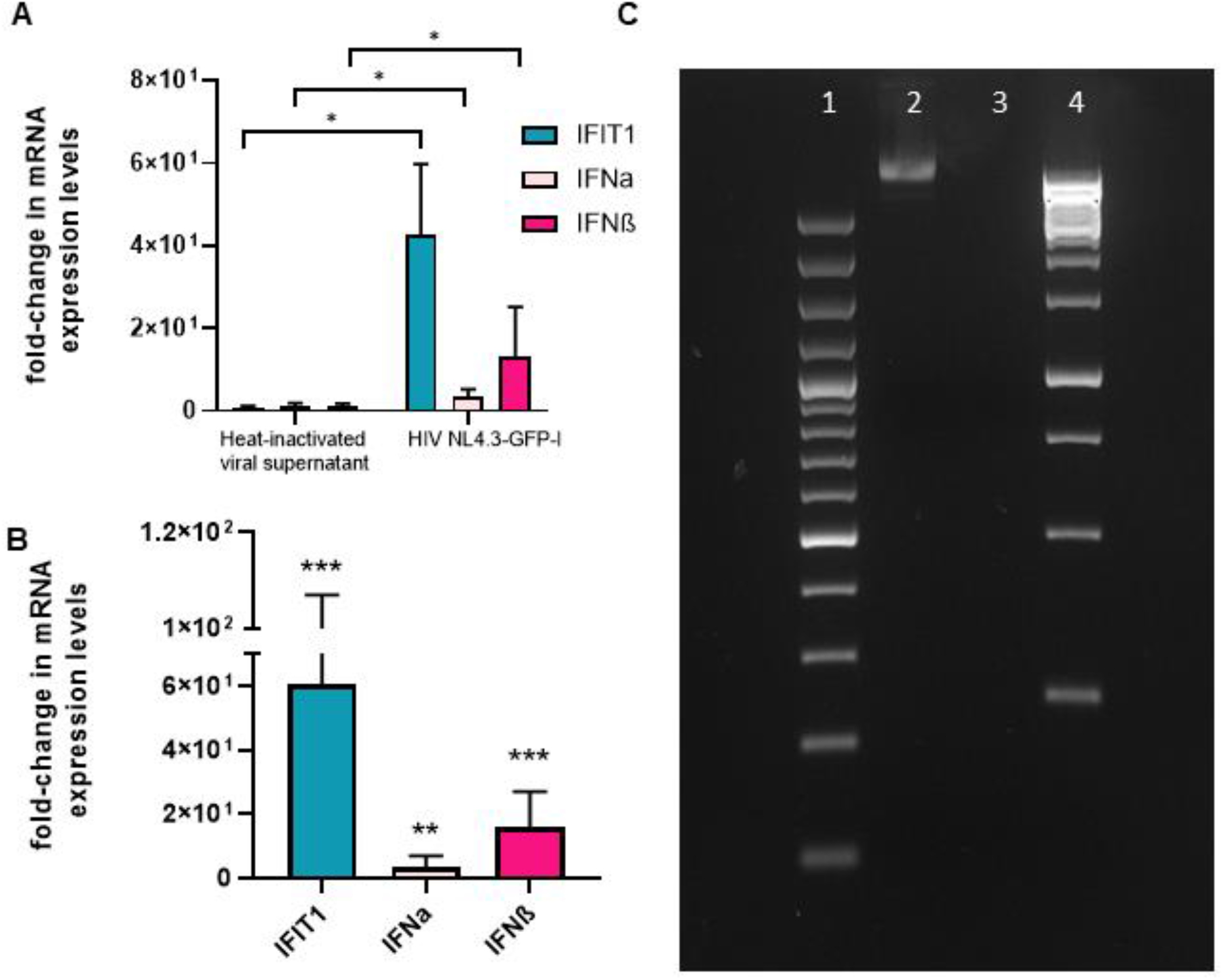
Plasmid contamination is not responsible for innate response of CD4+ T cells to HIV infection. **A** Heat inactivation of virus eliminates innate response. HIV-1 NL4-3-GFP-I virus was produced in a 293T cell line following transfection with a molecular clone as described before [10], and propagated in activated primary human PBMC cells. The supernatant was used for further production in a Jurkat CD4 CCR5 cell line and finally collected. The viral sup underwent heat-inactivation at 95 °C for 30 minutes. Figures show fold-change in mRNA expression levels for IFIT1, IFNα and IFNβ compared to non-infected cells with or without heat inactivation. Data shown are accumulative from 6 different experiments (n = 6 donors) mean ± SD * p<0.0313 **B** DNAse treatment of viral supernatant does not eliminates innate response. Plots show fold-change in mRNA expression levels for IFIT1, IFNα and IFNβ between cells infected with benzonase-treated viral supernatant of HIV-1 Nl4.3 GFP-I and uninfected cells. Data shown are accumulative from 11 different experiments (n = 11 donors) mean ± SD ***p<0.001; *** p<0.0059; *** p<0.001. **C** Activity of benzonase (Merck KGaA, Darmstadt, Germany) conform protocol by Sastry et al (21), 2004. Agarose gel with molecular weight markers (lanes 1,4) untreated plasmid (lane 2) and benzonase-treated plasmid (lane 3). For graphs **A** and **B** Wilcoxon matched pairs test was performed.

Next, HIV NL4-3-GFP-I viral supernatant was cleared from possible plasmids remnants by a treatment with endonuclease, relying on the protocol of Sastry et al, 2004 [12] for plasmid DNA removal (Fig. 1C). Endonuclease digested supernatant still clearly induced a response (Fig. 1B).

Finally, we propagated HIV-1 BaL [13], in addition to HIV-1 A8 and HIV-1 D2 strains [10] on activated primary human PBMC cells, and on susceptible cell lines, SupT1 (Fig S1 A), THP1 and Jurkat CD4 CCR5 (Fig S1 B) cells, respectively. Importantly, HIV-1 NL4-3-GFP-I was also propagated in this way to dilute out possible plasmid remnants; HIV-1 A8 and D2 are primary isolates not derived from plasmids. The CCR5-tropic HIV-1 BaL strain failed to infect the T cells at significant levels over a period of 7 days as expected, and consequently hardly any sensing was observed (Fig 3 A-C), which is in line with the observations by Elsner et al [11]. On the contrary, productive infection of the T cells with the HIV-1 D2 strain and cell-grown HIV-1 NL4-3-GFP-I induced high expression levels of IFIT1, IFNα and IFNβ, 4 days post-infection (Fig. 2 A-C). This induction was dependent on expression of cGAS (data not shown, confirming previous report [10]).

**Fig. 2.**
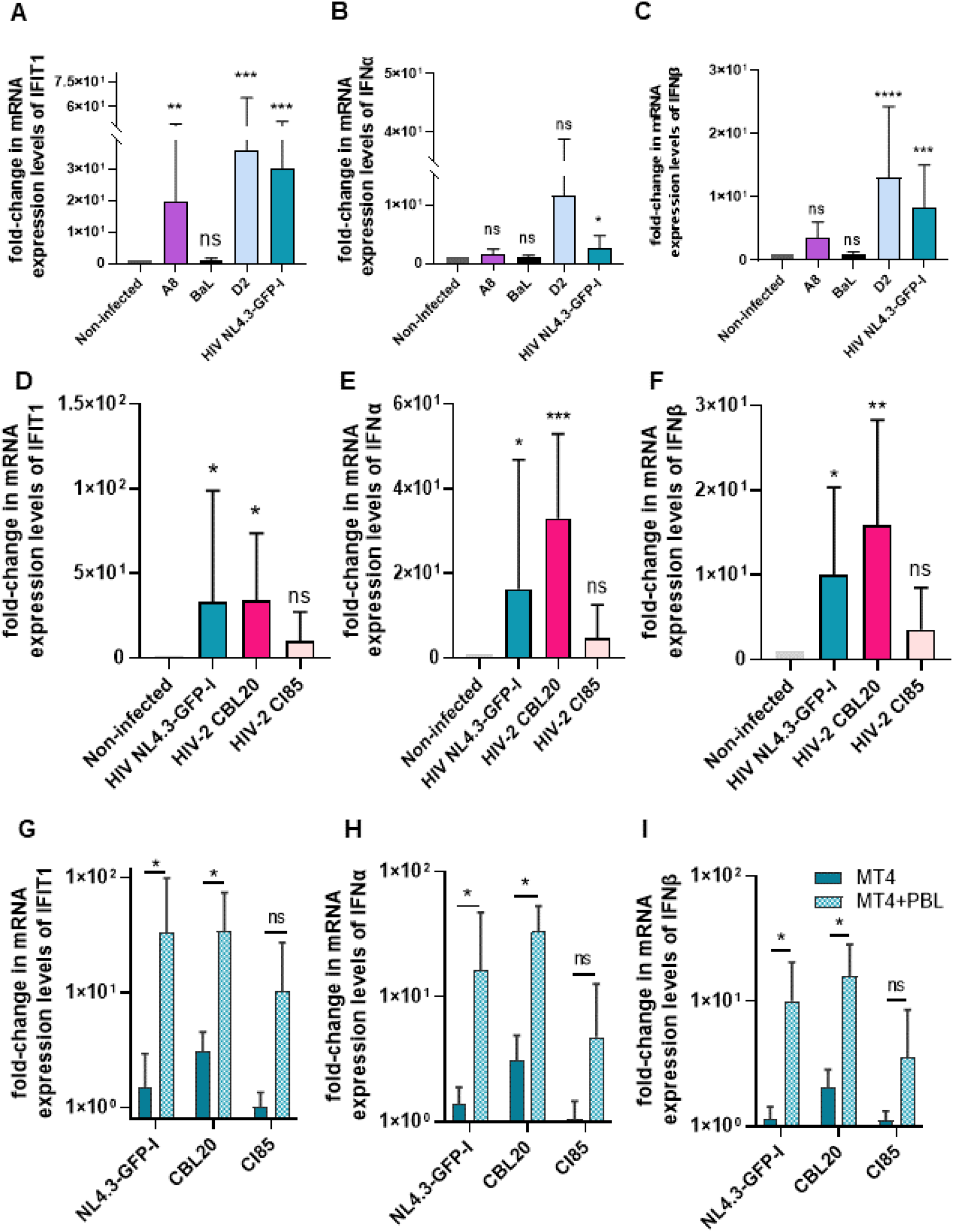
Induction of Interferon response after infection with HIV-1 and HIV-2. **A-C** Induction of IFN-I response by different HIV-1 viruses in infected primary T cells. Plots show fold-difference of mRNA expression levels compared to non-infected controls (n=10 for A8, BaL, D2 and HIV-1 NL4-3-GFP-I). IFIT1 A8 ** p<0.007, BaL ns p<0.6858, D2 *** p<0.0003, HIV NL4.3-GFP-I *** p<0.0005; IFNα A8 ns p<0.1198, BaL ns p<0.999, D2 ns p<0.3222, HIV NL4.3-GFP-I * p<0.0162; IFNβ A8 ns p<0.0591, BaL ns p<0.6858, D2 **** p<0.0001, HIV NL4.3-GFP-I *** p<0.0005. **D-F** When compared, both HIV-1 and HIV-2 induce a strong IFN-I response, however the amount of IFN induced seems to be dependent on HIV subtype. Data shows fold-difference of mRNA expression levels of IFIT1, IFNα and IFNβ compared to non-infected controls. IFIT1 HIV NL4.3-GFP-I * p<0.0389, HIV-2 CBL20* p<0.0214, HIV-2 CI85 ns p<0.9019; IFNα HIV NL4.3-GFP-I * p<0.0214, HIV-2 CBL20 *** p<0.0003, HIV-2 CI85 ns p<0.6426; IFNβ HIV NL4.3-GFP-I * p<0.0113, HIV-2 CBL20 ** p<0.0028, HIV-2 CI85 ns p<0.999. **G-I** HIV infection of MT4 cells does not induce a significant IFN-I response compared to infected CD4+ T-cells in co-culture. Data shows difference in fold-change in gene expression levels of IFN stimulated genes in MT4 cells infected with HIV-1 or HIV-2 virus compared to fold-change of ISGs in co-culture of MT4 cells and CD4+ T cell. IFIT1 HIV NL4.3-GFP-I * p<0.0313, HIV-2 CBL20 * p<0.0313, HIV-2 CI85 ns p<0.0781; IFNα HIV NL4.3-GFP-I * p<0.0313, HIV-2 CBL20 * p<0.0313, HIV-2 CI85 ns p<0.2188; IFNβ HIV NL4.3-GFP-I * p<0.0156, HIV-2 CBL20 ** p<0.0313, HIV-2 CI85 ns p<0.0781. Graphs show data ± mean. For graphs A-F Friedman’s test followed by Dunn’s multiple comparison test was performed, comparing infected cells to non-infected controls. For graphs G-I Wilcoxon matched pairs test was performed, comparing MT4 cells to the co-culture.

**Fig. 3.**
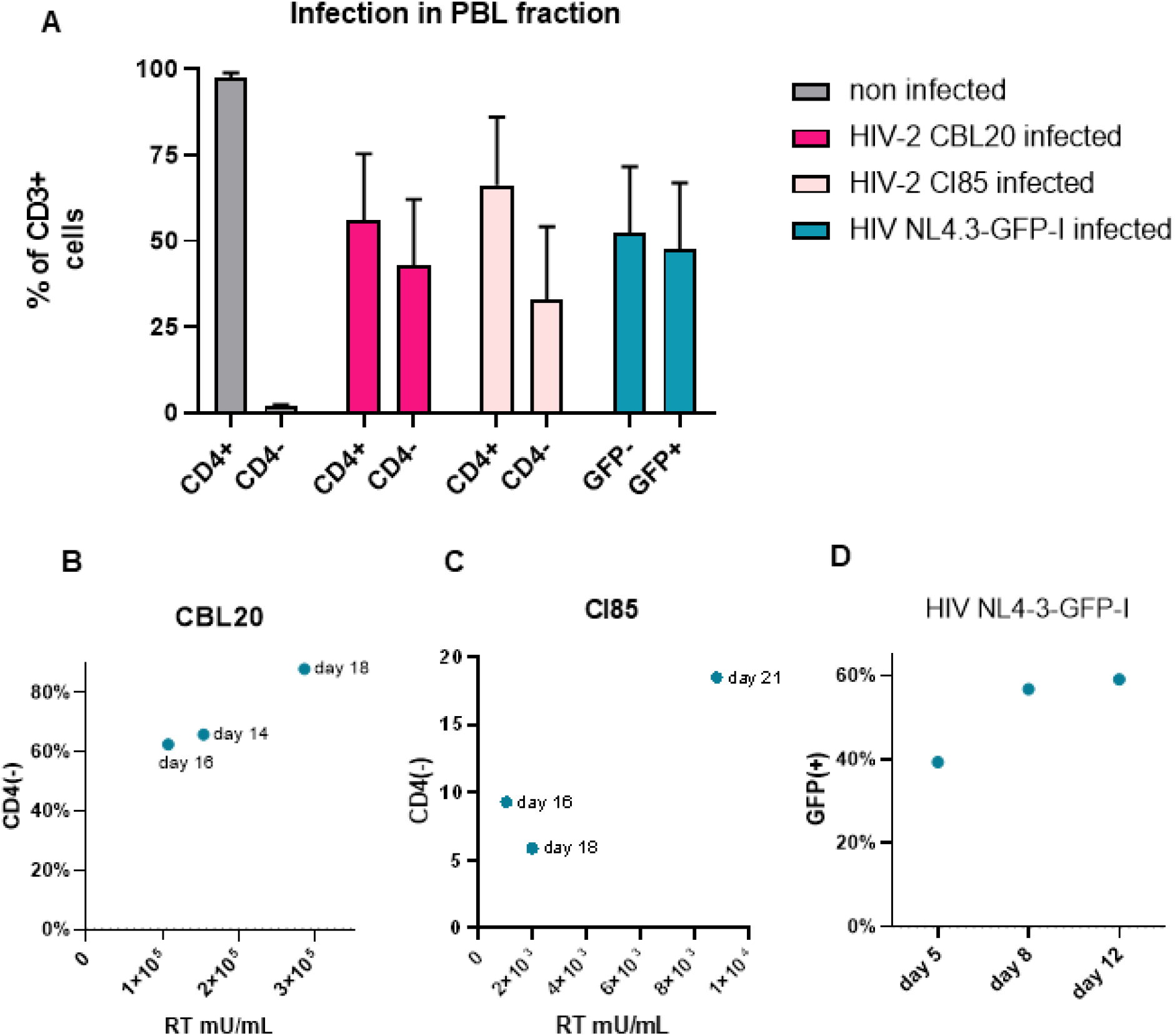
Monitoring rate of HIV-1 and HIV-2 infection. **A** Plot shows assessment of HIV-1 and 2 infections in CD4+ T cell fraction in co-culture with MT4 cells. Data analyzed after flow cytometry by gating on the CD3+ fraction (primary T cells) and subsequently gating the CD4+CD3+ fraction (non-infected cells) and CD4-CD3+ fraction (infected cells). Infection is clearly shown as a CD4-CD3+ fraction of the co-culture. Non-infected cells show almost no CD4-CD3+ fraction present, as expected. HIV-1 NL4-3-GFP-I infection can be also assessed using the GFP marker. Data is analyzed using the MACS Quantify software. **B-C** Infection with HIV-2 strains can be monitored by assessing the down-regulation of the CD4 marker in infected cells. Plots shows correlation of CD4 negative fraction with calculated RT values in viral supernatant harvested at that moment. To perform absolute quantification of RT activity values, a standard curve of replication-competent HIV-1 containing supernatant with known RT activity levels was run in parallel in each assay and values were extrapolated from the obtained Cq values. **D** Similarly, infection with HIV-1 NL4.3-GFP-I can be monitored with the GFP marker in infected cells. Plot shows increase of GFP+ cells per day to show progression of infection.

### Both HIV-1 and HIV-2 strains which effectively infect CD4+ T cells induce an innate response

In order to determine if several strains of HIV-2 induce IFN-I response and change the expression levels of IFIT1, IFNα and IFNβ, in the same way as HIV-1 infection [10], we compared primary isolates of HIV-2 (CBL20 and CI85) with HIV-1 NL4-3-GFP-I virus in infecting PHA/IL-2 activated primary CD4+ T cells by co-culturing them with infected MT-4 cells. As we intend to study primary isolates of HIV-2, we cannot rely solely on marker gene expression or p24 staining, respectively. Therefore, we first validated CD4 downregulation in these primary T cells as a marker for infection. Supplemental figure 2 shows flow cytometric analysis on T cell lines, demonstrating CD4 downregulation by both HIV-2 isolates, compared to HIV-1 NL4-3-GFP-I virus. In primary CD4 T cells, the fraction of CD4-cells increased with rising RT values in corresponding cell culture supernatants harvested at the same time (Fig. 3 B-D). Infection with both HIV-1 (Fig. 2 A-C) and HIV-2 (Fig. 2 D-F) showed comparable responses as measured in the change of gene expression levels of three different interferon-stimulated genes, IFIT1, IFNα and IFNβ. As a control, HIV-1 or HIV-2 infected MT4 cells alone do not induce IFN-I response (Fig 2 G-I), proving that the interferon response we detect after infection originates from the primary activated CD4+ T-cells.

In all experiments, primary CD4+ T cells were thoroughly depleted of plasmacytoid dendritic cells prior to infection, excluding the latter as the source of IFN-I (Figure S3). The levels of HIV-induced IFN-I varied in CD4+ T cells derived from different donors, but usually progressed in the same manner upon infection. Similarly IFN-I response correlated with the amount of spreading infection (percentage of CD4-cells) with the different primary isolates of HIV-2 and HIV-1 NL4-3-GFP-I (data not shown). HIV subtypes that showed less productive infection, also showed proportionally smaller interferon response, as expected.

Altogether, these results demonstrate that HIV-2 infection is capable of eliciting a comparable innate response in quantitatively and timely comparable manner as HIV-1 infection.

### cGAS is essential for triggering immune response in CD4+ T cells for both HIV-1 and HIV-2 infection

Since it’s discovery in 2013 [6], cGAS has been regarded as a pivotal component in the host’s innate immune responses to viral infections, including in the case of HIV-1. Initial knock-down screening and further experiments have consistently demonstrated the significance of cGAS in orchestrating IFN-I production upon HIV-1 infection [10]. In this paper, we extended the understanding of the role of cGAS in the innate immune response to viral infections, specifically focusing on its involvement in the context of HIV-2 infection. Building upon its recognized significance in the host’s defense against viral invaders, particularly HIV-1, our study elucidates whether cGAS plays a similarly crucial role in sensing and responding to HIV-2.

We employed cGAS (MD21D1) knockdown strategies using with cGAS shRNA-encoding lentiviral vectors in primary CD4+ T cells. Our experimental design allowed us to assess the impact of cGAS knockdown on the expression levels of key immune response genes—IFNα, IFNβ, and IFIT1—following infection with both HIV-1 and HIV-2 (Fig 4). Despite inherent variations among individual donors, our results consistently demonstrated a substantial decrease in the induction of type I interferons (IFN-I) upon cGAS knockdown in primary CD4+ T cells.

**Fig. 2.**
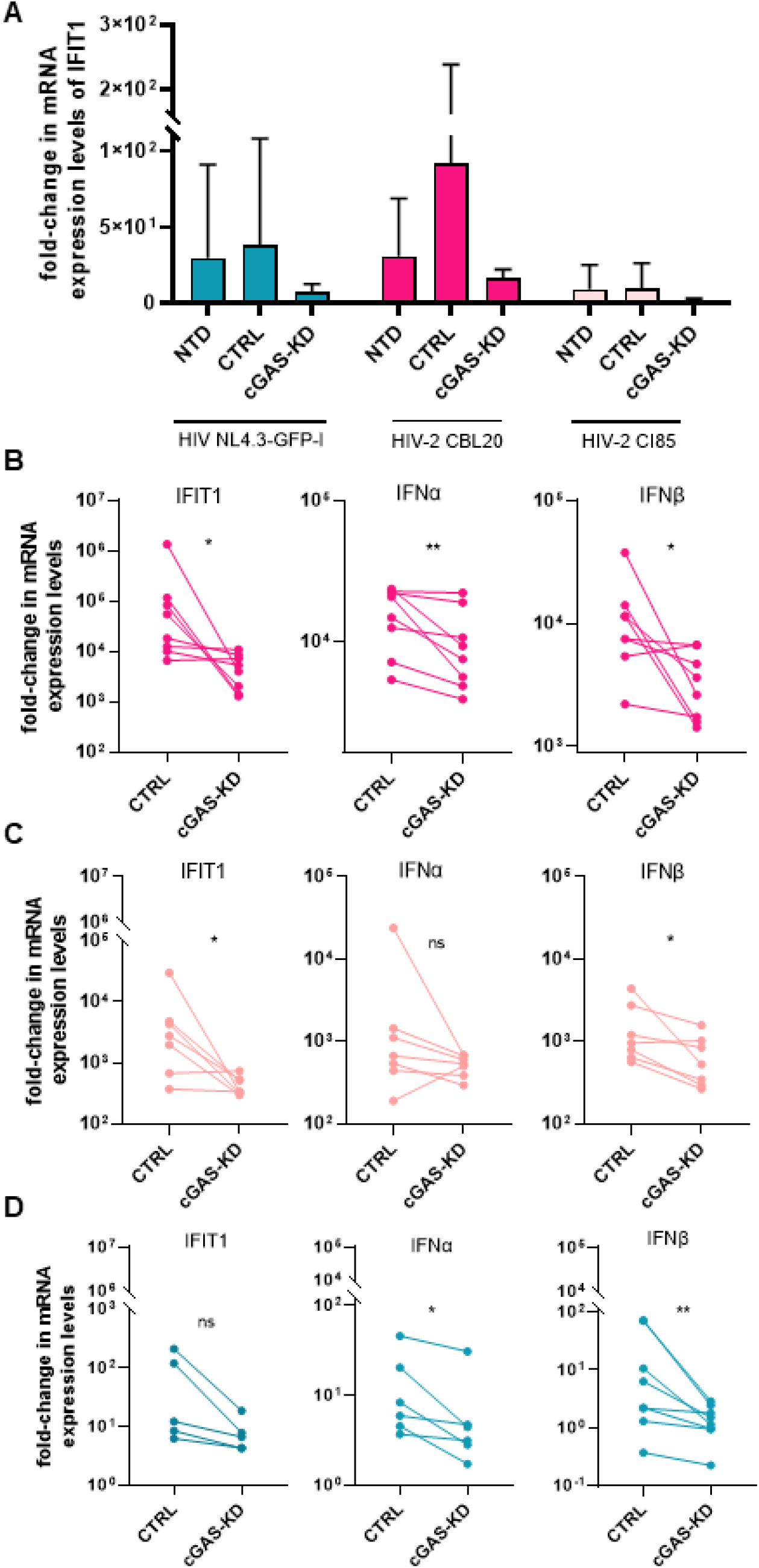
Knockdown of cGAS shows elimination of IFN-I response after infection with HIV-1 and HIV-2 virus. **A** Plot shows fold-difference in mRNA expression levels of IFIT1 after infection with different HIV strains in non-manipulated (non-transduced cells, NTD), control cells transduced with scr-puro control lentiviral vector (CTRL) and cells transduced with shRNA-encoding lentiviral vector against cGAS (cGAS). **B-D** Plots show individual data points of transduced cultures used for calculating average shown in A. Fold difference in mRNA expression levels of IFIT1, IFNα and IFNβ in cells knockdown for cGAS after infection with HIV-2 CBL20 (Panel **B**: IFIT1 * p<0.0156 ; IFNα ** p<0.0078 ; IFNβ * p<0.0391), HIV-2 CI85 (Panel **C**: IFIT1 * p<0.0469 ; IFNα ns p<0.1094 ; IFNβ * p<0.0313); HIV-1 NL4.3-GFP-I (Panel **D**: IFIT1 ns p<0.0625 ; IFNα * p<0.0313 ; IFNβ * p<0.0078) is shown. Wilcoxon matched pairs test was performed.

The comparative analysis between HIV-1 and HIV-2 infections provided insightful findings, highlighting the conserved nature of the cGAS-dependent type I interferon response in CD4+ T cells across both HIV types.

## Discussion

Innate sensing mechanisms play a crucial role in controlling viral infections by triggering interferon (IFN) responses. Over the years, the understanding has deepened, revealing that viruses, including HIV, are sensed by various cells. However, HIV-1 has evolved various strategies to evade immune detection and establish persistent infection, contributing to the challenges in developing an effective immune response against the virus. By studying the innate sensing of HIV-2 we might understand more the differences in pathogenicity between the two viruses [14,15] and understand natural control of infection. This lesser-known variant of the virus has been comparatively less studied, but presents a unique opportunity to gain insights into the spectrum of HIV pathogenicity.

The Herpesviridae family have been shown to trigger a cGAS-dependent antiviral response upon infection. By initially sensing the viral dsDNA, the DNA sensor cGAS induces intracellular defense response culminating with expression of type I interferons (IFNs). Accumulating data on cGAS-triggered immune responses [7,10,16,17]. in HIV-1 and HIV-2 infections prompted a comparative analysis, particularly considering the slower progression and long-term asymptomatic period in HIV-2. This led us to investigate the innate sensing of both types in activated CD4+ T cells, shedding light on potential differences in their responses.

Addressing concerns raised by Elsner et al. [11] about plasmid contamination in virus preparations triggering this response, we meticulously investigated this. Our previous work [10] had shown that activated CD4+ T cells sense HIV-1 infection through cGAS and induce an IFN-I response. Some but not all viral preps used in that work were started from proviral plasmids (like HIV-1 NL4-3 GFP-I virus). Elsner et al. [11] suggested that HIV-1 is not recognized by cGAS in infected T cells, however they used a monotropic HIV-1 strain BaL [13] which hardly infects T cells as we confirm here (Fig 2 A-C and S1). Moreover, previous findings by Vermeire et al. [10] demonstrated that the induction of interferon (IFN) was significantly reduced when viruses lacking integrase or tat proteins derived from proviral plasmids were exposed to activated CD4+ T cells, or when treating plasmid-produced HIV-1 NL4-3 GFP-I infected T cells with a reverse transcriptase or integrase inhibitor. This strongly argues in favor of the need for productive infection and against involvement of plasmid contamination in the sensing observed. Furthermore, as we show here experiments using plasmid-free viral preparations produced by viral replication on cell lines or PBMCs, such as two CXCR4-tropic primary HIV-1 group M strains (A8, D2) (Fig. 2 A-C) and an HIV-2 isolates CBL20 and CI85 (Fig. 2 D-F), also disprove the claim that sensing of plasmids explains the innate response we observe after HIV infection.

We employed a number of experimental approaches to further address this concern, and our findings collectively showed that the innate response of CD4+ T cells to HIV infection is not an artefact caused by contaminating plasmid since it is heat-sensitive and endonuclease treatment resistant. In addition, HIV NL4-3-GFP-I was also propagated on cell lines to dilute out possible plasmid remnants. Again, upon productive HIV-1 infection innate sensing was observed, dependent on expression of cGAS (data not shown, confirming previous report[10]). Innate immune response upon wild-type HIV-2 infection of non-manipulated primary cells has thus far been observed in primary plasmacytoid and myeloid dendritic cells[7–9], and in activated CD4+ T cells[10]. We extend previous observation here with the primary HIV-2 isolate CBL20 that induces a strong IFN-I response, at least as strong as in HIV-1 infection. However, the CI85 HIV-2 isolate was less potent. Therefore, the difference in progression of infection between the two HIV types could not be definitely attributed to a difference in quality of innate sensing as such. A recently published study shows that HIV-2 infection induces a stronger innate immune response in macrophages, compared to HIV-1 (group M) infection [17]. In our experiments, a significant contribution of myeloid cells to the IFN-I response measured is unlikely, given the purification of CD4+ T cells (fraction of remaining CD123+CD304+CD303+ pDCs in all of our experiments was below 0.01%, (Figure S3)).

Similar to HIV-1 infection, our results show the essential role of the DNA sensor cGAS in sensing of HIV-2 infection in activated CD4+ T cells. In the past there were questions about how and if HIV DNA is being sensed in the cytosol, due to the known fact that the mere production of HIV DNA during infection is not sufficient for immune activation. However, Lahaye et al. [7] recently demonstrated that despite cGAS being a sensor primarily located in the cytosol, it actually detects HIV DNA within the nucleus through its interaction with the capsid sensor NONO which interestingly enough appears to have a stronger affinity for the HIV-2 capsid [16,17]. This is consistent with HIV-2’s lower pathogenicity and superior immune control. This study noted increased HIV-1 infection post-NONO knockdown in Jurkat and primary CD4+ T cells, but its impact on immune response and sensing is unexplored. Additionally, reversion of HIV-1 capsid mutations absent in HIV-2 appears to be enhancing cGAS sensing in monocyte-derived macrophages (MDM) [17], triggering a stronger IFN-I response compared to wild-type virus. This IFN-I response was dramatically reduced by cGAS depletion. Distinctively, in our study we show that upon cGAS knockdown in the primary target of HIV-1 and 2, CD4+ T cells, we see clear reduction of expression of IFIT1, IFNα and IFNβ upon HIV-1 and HIV-2 infection, confirming the importance of cGAS as a crucial sensor involved in sensing of HIV infection in both macrophages and CD4+ T cells.

We previously showed that IFN-I response in activated CD4+ T cells is only induced after integration of the virus and expression of new viral proteins, since integrase- or tat-deficient viruses or viruses cultured with an integrase inhibitor induced insignificant or strongly reduced levels of IFN-I [10]. This indicates that sensing of HIV DNA is enabled through the action of newly expressed HIV replication products. In addition, in MDM, Lahaye et al [16] showed that HIV capsid can be sensed in the nucleus in infected cells, using an integrase inhibitor to restrict incoming viral capsids, however they don’t show if this sensing results in productive IFN-I response. Whether despite sensing, cGAS-mediated IFN response seems to be happening only post-integration, in MDM as we previously observed [10] in activated CD4+ T cell remains to be answered.

Our work ultimately demonstrates that HIV-1 and HIV-2 sensing in its main target cells, CD4+ T cells, depends on productive infection and cGAS. Since the magnitude differs between isolates regardless the type, the difference in progression of infection between the two HIV types can at present not be explained by intrinsic difference in innate sensing in activated CD4+ T cells.

## Materials and methods

### Production of shRNA lentiviruses

Lentiviral vector production from pLKO.1 vectors in 293T cells was done as reported previously[18]. 293T cells were seeded 24 h before transfection in 6-well plates at density of 400,000 cells per well. Separate co-transfections of the shRNA pLKO.1 clone targeting cGAS (pLKO.1 puro-shRNA against MB21D, TRCN0000148694) and a control scrambled shRNA pLKO.1 clone (SHC-002) were performed using MISSION Lentiviral Packaging Mix (Sigma-Aldrich, Diegem, Belgium) and FuGENE HD (Promega, Madison, WI) per manufacturer’s instructions. The supernatant was harvested 48 h and 72 h after transfection and stored in small aliquots at -80 °C until further use.

### Determination of lentiviral titer

The lentiviral titer was measured by quantification of reverse transcriptase activity (RT) via real-time PCR and expressed as equivalent amount of p24 protein [19]. These lentiviral supernatants were found to express RT activity of 14,000 – 20,000 mU/mL (equivalent of 2.6 - 3.6 μg of p24/mL) and were used in ratio 250 mU RT (equivalent to 45 ng p24) of virus to 1.5 × 10^6^ cells per well in 6-well plate.

### HIV viruses

The CXCR4-tropic HIV-1 group M A8 and D2 strains, as well as the primary HIV-2 CI85 and CBL20 isolates were kindly donated by the AIDS clinic at the Institute of Tropical Medicine in Antwerp, Belgium, with the approval of the ethics committee after written informed consent.

CCR5-tropic HIV-1 group M Ba-L strain was obtained through the NIH HIV Reagent Program, Division of AIDS, NIAID, NIH: Human Immunodeficiency Virus 1 (HIV-1) WITO4160.27 Env Optimized Expression Vector (pWITO4160.27 gp160-opt), ARP-11408, contributed by Drs. Beatrice H. Hahn, Jesus Salazar-Gonzalez, Denise L. Kothetr.

The replication-competent HIV-1 virus HIV NL4-3-GFP-I virus was kindly donated by Dr D.N. Levy [20,21]. Viral stocks were initially obtained after propagation in short-term cultures of PBMCs and expanded by infection and culture of Jurkat CD4 CCR5 cells. Supernatant of the Jurkat CD4 CCR5 and THP1 cells was collected at peak of infection, depending on viral kinetics after 18 to 29 days.

### Elimination of plasmid DNA in viral supernatants by endonucleases

To reduce possible plasmid contamination in HIV-1 NL4-3-GFP-I supernatant, in some experiments the supernatant was treated with benzonase (Merck KGaA, Darmstadt, Germany), as described by Sastry et al [12].

### Heat-inactivation of viruses

HIV-1 NL4-3-GFP-I supernatant was heat-inactivated to demonstrate no heat-resistant plasmids and other components could account for innate sensing. For this, tubes with the collected supernatant were incubated at 95°C for 30 minutes, eliminating infectious virus.

### Isolation of primary CD4+ T cells

Primary CD4+ T cells were isolated from buffy coats of healthy donors (Red Cross, Ghent, Belgium), donated after informed consent, approved by Ghent University Hospital ethical committee. Peripheral blood mononuclear cells (PBMCs) were obtained after Lymphoprep (Axis-Shield PoC, Oslo, Norway) centrifugation and used for CD4+ T cell isolation by negative selection with a commercial kit (Miltenyi Biotec, Bergisch Gladbach, Germany) per manufacturer’s instructions. Depletion of plasmacytoid dendritic cells (pDCs) was done on PBMCs prior to isolation of CD4+ T cells, by negative selection after staining with CD304-PE antibody (clone AD5-17F6, Miltenyi Biotec) in the presence of human FcR blocking reagent (Miltenyi Biotec) and subsequent staining with anti-PE paramagnetic microbeads (Miltenyi Biotec) as shown before [10]. Purity of CD4+ T cell population was measured by flow cytometry (MACSquant Analyzer using MACSQuantify software, Miltenyi Biotec), showing a fraction of at least 95% CD4+CD3+ double positive cells and less than 0.01% CD123+CD304+CD303+ cells. The assessed purity was always found to be above 95%.

Except when used for lentiviral transduction (see below), the fresly isolated primary CD4+ T-cells were cultured in 96-well plates at 300,000 cells/well in 0.1 mL RPMI medium (RPMI 1640, Life Technologies, Carlsbad, CA) supplemented with 2 mM L-glutamine (Life Technologies), 10% (v/v) heat inactivated fetal calf serum (FCS, Hyclone, Thermo Scientific, Rockford, IL), 100 U/mL penicillin, 100 μg/mL streptomycin (Life Technologies), interleukin-2 (IL-2; specific activity 10 U/ng) at concentration of 20 ng/mL (Roche) and phytohemagglutinin (PHA) mitogen at 1 μg/mL (Sigma-Aldrich, Remel) at 37°C in a humidified atmosphere containing 5% CO_2_.

### shRNA-mediated gene knock-down in primary CD4+ T cells

Primary CD4+ T-cells depleted of plasmacytoid dendritic cells were immediately transduced with pLKO.1 lentiviral vectors upon isolation from peripheral blood. The cells were subsequently plated at a density of 300,000 cells in 55 μL RPMIc per well of a 96-well flat-bottom plate. Lentiviral vector supernatant (45 μL) was then added to each well in the presence of interleukin-2 (20 ng/mL), PHA (1 μg/mL) and polybrene (8 μg/mL; Sigma-Aldrich). Three different conditions were employed, namely: 1) cells transduced with lentiviral vector encoding a scrambled shRNA sequence (Non-targeting scrambled shRNA control (SHC-002)), 2) cells transduced with lentiviral vector encoding shRNA sequence that targets cGAS and 3) non-transduced cells.

Cells were subsequently spinoculated (30 min, 950 g, 32 °C) in the presence of polybrene (8 μg/mL). 48 hours after transduction, puromycin (1.2 μg/mL final concentration, Sigma-Aldrich) was added to the cells. After 72 h, PHA stimulation was stopped by removing all medium and adding fresh RPMIc containing IL-2 and puromycin. (10)

### HIV infection

#### Primary CD4+ T cells

After isolation, cells were infected with different primary isolates of HIV-1 (BaL, A8, D2) as well as HIV-1 NL4.3-GFP-I virus by plating 1.5 × 106 cells per well of a 6-well flat-bottom plate (BD Bioscience, X, Y) and adding 250 mU RT (equivalent to 45 ng p24) of virus in a total volume of 6 mL IMDMc. Cells were subsequently spinoculated at 2300 rpm for 90 minutes at 32 °C.

#### Co-culture of MT4 cells with primary CD4+ T cells

At day 4 after shRNA-mediated gene knock-down, parallel cultures of MT4 cells (between passage 3 and 7) were infected with two different primary isolates of HIV-2 (CI-85 and CBL20) and HIV-1 NL4-3-GFP-I virus by plating 1.5 × 106 cells per well of a 6-well flat-bottom plate (BD Bioscience, X, Y) and adding 250 mU RT (equivalent to 45 ng p24) of virus in a total volume of 6 mL IMDMc. Cells were subsequently spinoculated at 2300 rpm for 90 minutes at 32 °C. 48 h after infection, cells were collected and used for co-culture with primary CD4+ T-cells. HIV-1 and HIV-2 infection was measured using flow cytometry gating on CD4-CD3+ cells as judged by CD4 down-regulation [22,23] of CD4+ T cells upon infection or by the GFP marker in case of HIV NL4-3-GFP-I virus.

Using a co-culture system for indirect HIV infection allowed for efficient infection of primary cells within 24 h[10]. This system was chosen to avoid the use of long-term cultures of HIV-infected primary cells that are also lentivirally transduced, as the MT4 cell line (an HTLV-I-transformed T-cell line) notably supports the growth of HIV infection [24–26].

MT4 cells were cultured with complete IMDM medium (Life Technologies) supplemented with 10% (v/v) fetal calf serum, 2 mM L-glutamine, 100 U/mL penicillin and 100 μg/mL streptomycin at 37 °C in 7% CO_2_ (v/v) in humified atmosphere.

#### Antibodies and flow cytometry

Antibodies used for staining were CD3-PE (clone SK7, Becton Dickinson (BD) Biosciences, Erembodegem, Belgium) and CD4-APC (clone MT4 66, Miltenyi Biotec) or CD123-FITC (clone AC145, Miltenyi Biotec), CD303-APC (clone AC144, Miltenyi Biotec) and CD304-PE.

To check for infection rates of MT4 cells before co-culture with primary CD4+ T cells, MT4 cells were stained with CD4-APC antibody to measure the CD4 downregulated fraction. In primary CD4+ T cells, HIV-1 and HIV-2 infection was measured 24 hours post-infection using flow cytometry gating on CD4-CD3+ cells as judged by CD4 down-regulation of CD4+ T cells upon infection.

In co-cultures, MT4 cells were seen as CD4+CD3-cells, CD4+ T cells that were not infected were CD4+CD3+ double positive cells, while infected CD4+ T cells were seen as CD4-CD3+ T cells.

To analyze the data, both the MACS Quantify and the FlowJo software were used.

#### qPCR for evaluating mRNA expression after shRNA-mediated knock-down for cGAS

RNA was isolated from QIAZOL lysates using miRNeasy mini kit per manufacturer’s instructions. RNA (max 1 μg) was subsequently treated with amplification-grade DNAse I (Life Technologies) and used for synthesis of cDNA with Superscript III reverse transcriptase and random primers (Life Technologies). Depending on the gene to be measured, cDNA was subsequently diluted 3x (for target genes: IFNβ, IFNα and IFIT1) and 15x (for reference genes: ACTIN, RPL13A, YWHAZ and UBC) with Nuclease-free water (Ambion, Life Technologies). 5 μL of the diluted cDNA was then used for qPCR. For qPCR LightCycler 480 SYBR Green I Master mix (Roche Diagnostics, Vilvoorde, Belgium) was used in final reaction of 15 μL. qPCR reactions were performed in 384-well plates (LightCycler 480 Multiwell Plates 384, white, Roche Diagnostics) on the Light Cycler 480 II instrument (Roche Diagnostics).

All samples were measured in duplo. A non-template control (nuclease-free water instead of cDNA) and a serial 10-fold dilution of standard curve was used. The cDNA for the standard curve was synthesized using mRNA from poly(I:C) stimulated PBMCs and this standard curve was included for the measurement of each gene on the plate. Melting curve analysis for IFIT1, IFNα and IFNβ was performed and showed a single peak. Calibrated normalized relative quantities (CNRQs) were calculated for each target genes in each sample based on the obtained Cq values, with the qBase Software (Biogazelle, CellCarta, Montreal, Quebec), using YWHAZ, ACTIN, RPL13A and UBC as reference genes and using target- and run-specific amplification efficiencies.

#### Primers

Primers used for qPCR were: UBC Fwd (sense) 5’-ATTTGGGTCGCGGTTCTTG -3’, UBC Rev (antisense) 5’-TGCCTTGACATTCTCGATGGT-3’, YWHAZ Fwd (sense) 5’-CTTTTGGTACATTGTGGCTTC AA -3’, YWHAZ Rev (antisense) 5’-CCGCCAGGACAAACCAGTAT -3’, ACTIN Fwd (sense) 5’-TGACCCAGATCATGTTTGAGA -3’, ACTIN Rev (antisense) 5’-AGAGGCGTACAGGGATAGCA -3’, RPL13A Fwd (sense) 5’-CCTGGAGAAGAGGAAAGAGA -3’, RPL13A Rev (antisense) 5’-TTGAGGACCTCTGTGTATTTGTCAA -3’, IFIT1 Fwd (sense) 5’-GATCTCAGAGGAGCCTGGCTAA –3’, IFIT1 Rev (antisense) 5’-TGATCATCACCATTTGTACTCATGG -3’, IFNalpha Fwd (sense) 5’-GTGAGGAAATACTTCCAAAGAATCAC –3’, IFNalpha Rev (antisense) 5’-TCTCATGATTTCTGCTCTGACAA -3’, IFNbeta1 Fwd (sense) 5’-GCTTCTCCACTACAGCTCTTTC -3’, IFNbeta1 Rev (antisense) 5’-CAGTATTCAAGCCTCCCATTCA -3’. All primers were purchased from IDT (Integrated DNA Technologies, Europe Branch, Leuven, Belgium).

## Statistical analysis

Figures were created with Microsoft PowerPoint and GraphPad Prism version 8 for Windows (GraphPad Software, San Diego, California, USA). Statistical tests were performed with GraphPad Prism 8.0 as indicated in the figure legends.

## Data availability

Data will be shared upon reasonable request.

